# Characterizing Variants of Uncertain Drug Resistance (VUDRs) Using Quantitative Measurements at Clinical Exposures

**DOI:** 10.1101/2025.10.31.685392

**Authors:** Haider Inam, Marta Tomaszkiewicz, Joshua Reynolds, Zeyu Yang, Scott Leighow, Justin R. Pritchard

**Author notes:** Broad Institute of MIT and Harvard, MA 02142. Atlas Biotech, 200 Innovation Blvd, Suite 260A, State College, PA, 16803. These authors contributed equally (Haider Inam, Marta Tomaszkiewicz).

## Abstract

Somatic missense mutations in oncogenes drive resistance to anticancer drugs, yet many variants remain clinically uncharacterized. Analogous to Variants of Uncertain Significance (VUS) in genetic disorders, these Variants of Uncertain Drug Resistance (VUDRs) in cancer lack the functional annotation needed to guide clinical management. Here, we applied a standards-driven deep mutational scanning platform that connects quantitative concentration-response measurements to human pharmacokinetics across 4922 missense variants (>96% coverage) at approved (400 mg QD) and investigational (400 mg and 500 mg BID) human doses of imatinib. Resistance phenotypes for 18 standards spanning 2 orders of magnitude of drug sensitivity showed strong quantitative performance and clinical concordance. Analyzing 257 clinical VUDRs, >10% conferred modest levels of resistance that might be overcome by dose escalation with generic imatinib instead of a branded alternative, potentially alleviating financial toxicity. Integration with global germline data also revealed ancestry-specific variants with the potential to create private VUDRs. These preclinical data establish the first generalizable framework for high throughput resistance variant classification directly tied to known human doses.

## INTRODUCTION

### Variants of Uncertain Drug Resistance (VUDRs): Challenges in Drug Annotation at Therapeutic Doses and Limitations of Previous Drug Resistance Screens

Variants of Uncertain Significance (VUS) are mostly germline mutations commonly identified during genetic testing for hereditary disease risk, whose impact on disease predisposition is not yet known. Over the past decade, genomic sequencing has revealed that VUS are individually rare but collectively common, comprising 55.15% of over 3 million ClinVar entries^1–3^. This is a challenging medical question because the median frequency of a variant found in any one individual is as low as 1 in 1,000^4–6^, with protein-coding or functional variants being as rare as <1 in 10,000^7,8^. To address this critical challenge, high-throughput functional assays -- particularly multiplexed assays of variant effect (MAVEs), such as deep mutational scanning (DMS) or saturation genome editing (SGE) -- have been developed to systematically evaluate the functional effects of thousands of variants in parallel^9–11^. These approaches are particularly useful for disease-related genes, such as *BRCA1*^10,12,13^, *BRCA2*^14–16^, *TP53*^17^, and *PTEN*^18^, where accurate classification of rare missense variants is helpful for the diagnosis and clinical care of patients with a rare variant in a rare Mendelian inherited disorder.

In the case of *BRCA1* and *BRCA2*, DMS has enabled functional annotation of hundreds of missense variants, helping reclassify VUS as either likely pathogenic or benign. For example, studies such as Starita *et al.* 2018^10^ as well Findlay *et al.* 2018^19^ systematically evaluated *BRCA1* variants and demonstrated strong concordance between functional scores and ClinVar classifications. These efforts have informed clinical variant interpretation frameworks such as ABC and the ACMG/AMP guidelines^2,20,21^ and are increasingly cited in expert panel curation initiatives (e.g., by the ENIGMA consortium in their classification of *BRCA1*/*2* variants^22^). Importantly, Myriad Genetics, a major provider of hereditary cancer testing, incorporates both functional data and proprietary models into their variant classification algorithms^23^. Beyond *BRCA*, the NIH-funded Clinical Genome Resource (ClinGen) has endorsed the use of high-throughput functional evidence under specific criteria for variant interpretation, particularly when results are reproducible and validated^24^. These developments underscore a paradigm shift in functional genomics, moving from anecdotal evidence and low-throughput experiments to scalable, data-rich annotations that can keep pace with clinical sequencing.

Variants of Uncertain Drug Response (VUDRs), the term we introduce here, are typically somatic mutations arising in cancer in response to drug therapy with the potential to alter therapeutic sensitivity in humans at clinical doses. In cancer, genetic variation is acquired somatically during cancer initiation, progression, or treatment. These somatic variants arise on a background of standing germline variation and affect drug response^25–27^. Using the term VUDR is done to emphasize the distinct challenge of translating DMS findings into clinical predictions of drug response in light of known dosage forms and pharmacokinetics. Like VUS, VUDRs are rare individually but collectively more frequent across patients. However, unlike the growing array of tools developed for VUS clinical interpretation, current approaches for classifying VUDRs remain far from the clinical validation standards set by studies on *BRCA1*. A key reason for the evidence gap for VUDRs is that drug resistance is not a static genetic trait. Assessing drug resistance in the clinic relies on dose and concentration dependent Pharmacokinetic-Pharmacodynamic (PK/PD) relationships, e.g. different human doses create different concentrations of compound in vivo (PK) and different mutations have distinct concentration-response relationships (PD), thus adding complexity to high throughput screens and complicating interpretations of such data for drug resistance classifications^28–31^. Importantly clinically actionable relationships between individual in vitro assays and clinical exposures can be predictive of patient responses. This has led to the FDA approving and the NCCN recommending drugs for individual rare mutants on the basis of in vitro data alone, when no clinical data exists^32–34^. For instance, the label of ivacaftor^35^ shows that it is approved for 38 CFTR variants that were too rare to be included in the clinical trial, yet are thought to be sensitive to the drug. To take steps towards approving rare variants for on-label use, a high-throughput method would require extensive variant standards to demonstrate the reliability of a PK/PD relationship established between in vitro data and the patient responses at specific drug doses in people.

Recent attempts to use saturation variant profiling for drug resistance in cancer targets such as kinases: MAPK1/ERK2, CDK4, CDK6, HER2, MET, EGFR, and the GTPase KRAS^36–44^ have proven sufficient for the discovery of novel variants that impart a selective advantage in the presence of drug in vitro, and the variants discovered are enriched for resistance variants that are known to occur in the clinic and are useful. Existing studies also offer exciting insights into drug biology. However, all prior cancer drug resistance studies using high throughput techniques have utilized a single arbitrary drug concentration without quantitative standards that does not allow for the establishment of robust PK/PD relationships and comparisons to known clinical exposures at approved human doses. This means these insights cannot be used on an FDA label or in an NCCN guideline. Beyond this lack of concentration response data, and a lack of comparison to known levels in people, other barriers to quantitative assays and robust PK/PD relationships exist. These include: 1) the fact that the longer one selects for drug resistance the more the mean fitness of a pool of variants changes^45^. Metrics used to quantify resistance must be insensitive to this time-dependent change in pool fitness to avoid artifacts that can occur with the simple log2 fold change metrics that are pervasive in the drug resistance and genomics literature^45^, 2) the error rates (10^-2^-10^-3^) of conventional next-generation sequencing are too high for pool complexities that are much greater than 1000 variants, and while this has been discussed in the literature^46–48^ it has not been directly assessed as a factor limiting the quantitation of pooled drug resistance assays. To overcome some of these limitations, barcoding and error-correction sequencing (such as DMS-TileSeq or DMS-BarSeq) have been used to lower error rates in some deep mutational scanning (DMS) studies of Mendelian disorder genes^49^ but not cancer. However, the extent to which this error correction is necessary or sufficient for precise concentration response measurements and PK/PD relationships has not been evaluated.

Here, to address the challenges in predicting drug responses in VUDRs, we present a robust framework that applies error-corrected sequencing, growth-rate quantitation, clinical standards, and multiple concentrations to perform deep mutational scanning of imatinib resistance in BCR-ABL. Our approach overcomes key technical barriers to quantitation and accurately measures concentration-response profiles of clinically observed resistance mutations. Using these profiles we establish a clear PK/PD relationship across standards with a range of concentration-response profiles that allows for the interpretation of previously uncharacterized VUDRs at clinically relevant doses of imatinib, filling a major gap in the clinical annotation of drug resistance.

## RESULTS

### TileSeq balances sequencing error correction and measurement accuracy/sensitivity

A key challenge in deep mutational scanning (DMS) is that standard next-generation sequencing (NGS) is often too error-prone to accurately measure the lowest abundance variants being studied in complex pools^50–55^. The typical NGS error rate (10^-2^-10^-3^) is higher than the frequency of most variants in a typical DMS library (<10^-4^), making it difficult to distinguish a real signal from sequencing noise^47,48,56,50,51,57^. To confirm this, we re-analyzed a published DMS study on the *TP53* gene (a library comprising 8,258 *TP53* variants)^58^ and found that the read counts of a surprising number of true variants were indistinguishable from sequencing errors (Figure 1A). This result validates concerns that standard NGS can compromise the accuracy of DMS experiments.

**Figure 1.**
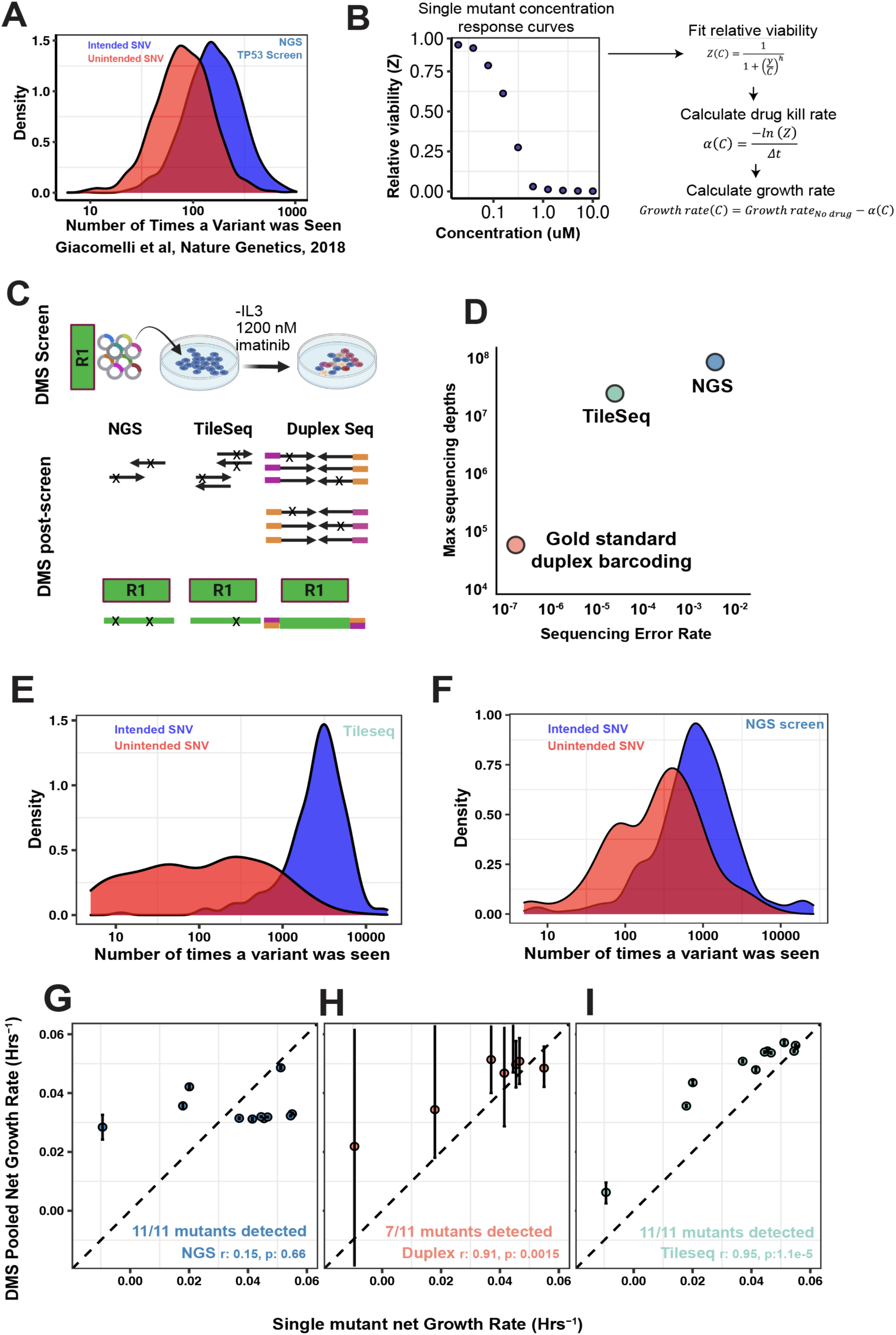
Trade-off between error rates and sequencing depth when quantifying resistance in DMS screens. **(A),** Count distribution of intended (blue) and unintended (red) single nucleotide variants detected in the TP53 DMS library^58^ (**B)** Single-mutant concentration-response curves and how drug-induced growth rates can be calculated using fits of these curves (**C)** Schematic of DMS screen conducted in BaF3 cells; region 1 refers to residues 242-321 of the BCRABL kinase (**D**), Sequencing error rate vs sequencing depths tradeoff across of three sequencing technologies: duplex sequencing (pink), Tileseq (cyan), and conventional NGS (blue) (**E, F**) Count distribution of intended (blue) and unintended (red) single nucleotide variants detected in region 1 of BCR-ABL by TileSeq and by NGS, (**G, H, I)** Correlation of net growth rates of mutant standards from single-mutant assays versus DMS measurements by NGS, Duplex Sequencing, and TileSeq. Mutants are a subset of the known clinically resistant mutants in region 1 of the BCRABL kinase. Error bars represent the binomial confidence intervals calculated using sequencing count data.

To accurately link lab measurements to clinical outcomes and potentially assess the impact of sequencing errors, we developed a panel of quantitative “gold standard” variants. This approach addresses a key limitation of previous DMS drug resistance studies, which lacked such standards and have not demonstrated accuracy and reproducibility^9,36–38,41,45,54,59^. For our *BCR-ABL* model, we leveraged 21 well-known mutations, individually measuring each across 10 drug concentrations to create precise concentration-response curves (Supplementary Figure 1, Supplementary Table S0). This panel provides a robust benchmark for quality control, spanning over an order of magnitude of sensitivity-from highly resistant mutants like T315I (IC_50_: 6123±1551 nM) to mutations like V299L (IC_50_: 185±50 nM) that is sensitive to imatinib but resistant to other ABL TKIs (Figure 1B). These standards form the foundation for our ultimate goal of building a quantitative clinical PK/in vitro PD relationship. Importantly, we also confirmed these standards perform reliably in pooled assays, provided that fitness is measured using a growth rate parameter (similar to a Chronos score^60^ or Enrich2^45^) instead of a static log₂ fold-change analysis (Supplementary Figure 2, Supplementary Table S1).

To address the high error rates of conventional NGS and the impact on gold standard measurements during a DMS screen we designed a pilot experiment to compare sequencing strategies. We focused on the N-lobe of the *BCR-ABL* kinase (residues 242-321; 1,517 substitutions), a region containing 11 of our 21 single-mutant standards. A site-saturation library was transduced into Ba/F3 cells at >500x coverage and screened for six days with 1200 nM imatinib (Figure 1C). We then assessed gold standard performance using 1) growth rate measurement correlations between pooled and unpooled data, 2) sensitivity to identify gold standard variants in the pool and 3) measurement error. Conventional NGS failed the primary validation test, yielding measurements that were completely uncorrelated with our unpooled single-mutant standards (Pearson’s r = 0.15) (Figure 1G). Duplex sequencing had an untra-low error rate of ∼1 in 10^7^, but suffered from critically low variant coverage, despite >100,000 unique molecules detected. With 90% of variants receiving only 1-10 reads (median = 3), it recovered just 7 of 11 standards (∼64%) and produced measurements with large confidence intervals (Fig. 1H, Supplementary Figs. 3C,D,E). TileSeq provided the optimal balance. It achieved deep coverage, with 90% of variants receiving 100 to 10,000 reads (median = 2,608), and detected all 11 out of 11 standards. Although TileSeq captures a higher proportion of unintended mutants than duplex sequencing (Supplementary Fig. 3A) it achieves markedly better separation between intended and unintended variants in the mutagenized region (Fig. 1E,F; Supplementary Fig. 3B), thereby improving true variant detection. Crucially, TileSeq’s measurements showed the highest correlation with the gold-standard data (Pearson’s r = 0.95) (Fig. 1I). These data reveal a critical trade-off between error correction and sequencing depth. Because TileSeq maintains high sensitivity while substantially reducing measurement error, we selected it as our preferred approach for all subsequent experiments.

### A High-Resolution Map at a Single Concentration of Imatinib Across the Full BCR-ABL Kinase Domain

Having established our method, we performed a DMS screen across the full-length *BCR-ABL* kinase domain. Using our TileSeq approach, we quantified the fitness of 4,925 single amino acid mutants at a high dose of 1200 nM imatinib (Fig 2A, Supplementary Table S2). The assay proved highly reproducible, with net growth rates showing a strong correlation between technical replicates (Pearson’s r = 0.92, p < 0.001, Fig 2B). To assess the accuracy of our pooled screen, we compared the results for 18 of our 21 standards that were recovered in the library (three were absent at residue 355). The pooled DMS growth rates correlated strongly with the unpooled single-mutant measurements (Pearson’s r = 0.8, p < 0.001), confirming our assay’s ability to precisely quantify resistance levels (Fig. 2C, Supplementary Fig. 2). To achieve a one-to-one concordance with our gold-standard data, we used these 18 standards to apply a linear regression correction (slope = 1.07 [95% CI: 0.64-1.49]; intercept = -0.02 [95% CI: -0.04-0.00]; Pearson’s r = 0.8) to the entire dataset. This standards-based correction eliminated a small systematic error and yielded a high-confidence imatinib fitness map for 4,925 variants, covering 96% of all possible single amino acid substitutions in the kinase domain. This resource enables the systematic interpretation of both rare and uncharacterized mutations at a single, clinically relevant drug concentration (Fig 2D, Supplementary Table S2).

**Figure 2.**
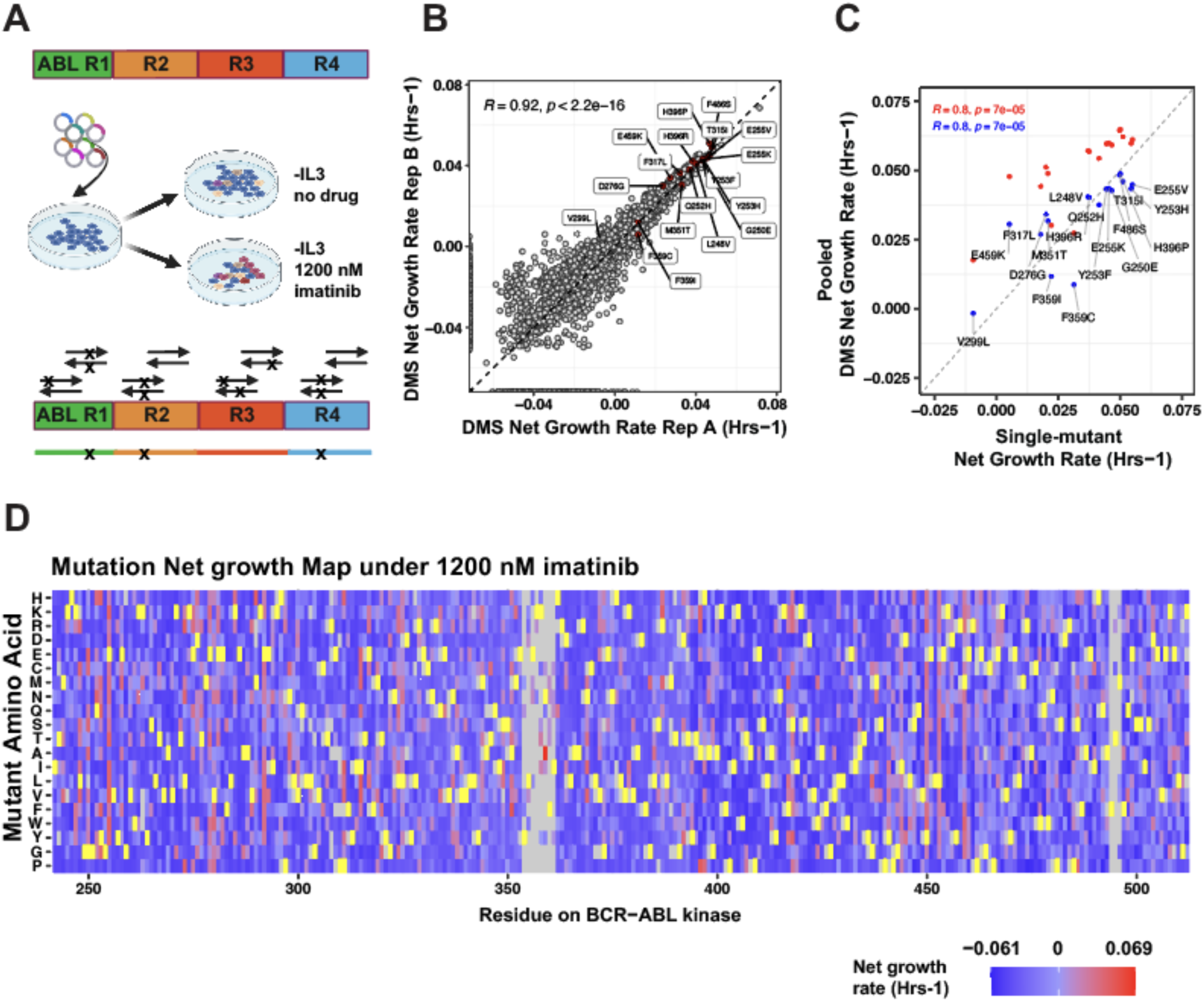
Validation and reproducibility of pooled DMS-based resistance profiling across the BCR-ABL kinase domain at a single concentration of imatinib. (**A**) DMS screen at an imatinib dose of 1200 nM across the full BCR-ABL kinase, (**B**) Correlation of net growth rates of 4,925 mutants across DMS replicates, Pearson’s r is shown (**C**) Correlation of net growth rates (red: without linear regression correction, blue: following correction) of mutant standards from single-mutant concentration response assays (x-axis) versus pooled-DMS measurements (y-axis), (**D**): Heatmap depicting net growth rates of 4,925 BCR-ABL mutants at 1200 nM imatinib in the DMS pooled assay. Yellow residues indicate WT amino acids. Grey residues indicate mutants missing from the cloning library.

### A Comprehensive Imatinib Concentration-Response Map for the Full BCR-ABL Kinase Domain

To move beyond a single concentration, we next sought to generate full concentration-response profiles for all variants. Since measuring 10 or more concentrations prohibitive for a DMS study, we first determined the minimal number of concentration points required. Using our standards, we found that a 3-point assay spanning a clinically relevant range (∼300nM to ∼1200nM) could reasonably recapitulate the absolute IC₅₀ values from a 10-point curve. For variants with IC₅₀ values below 1,200 nM, the correlation was nearly perfect (Pearson’s r = 0.99, p < 0.0001), though it deteriorated modestly for more resistant variants (Pearson’s r = 0.83, p = 0.02) (Fig 3A, Supplementary Table S6).

**Figure 3.**
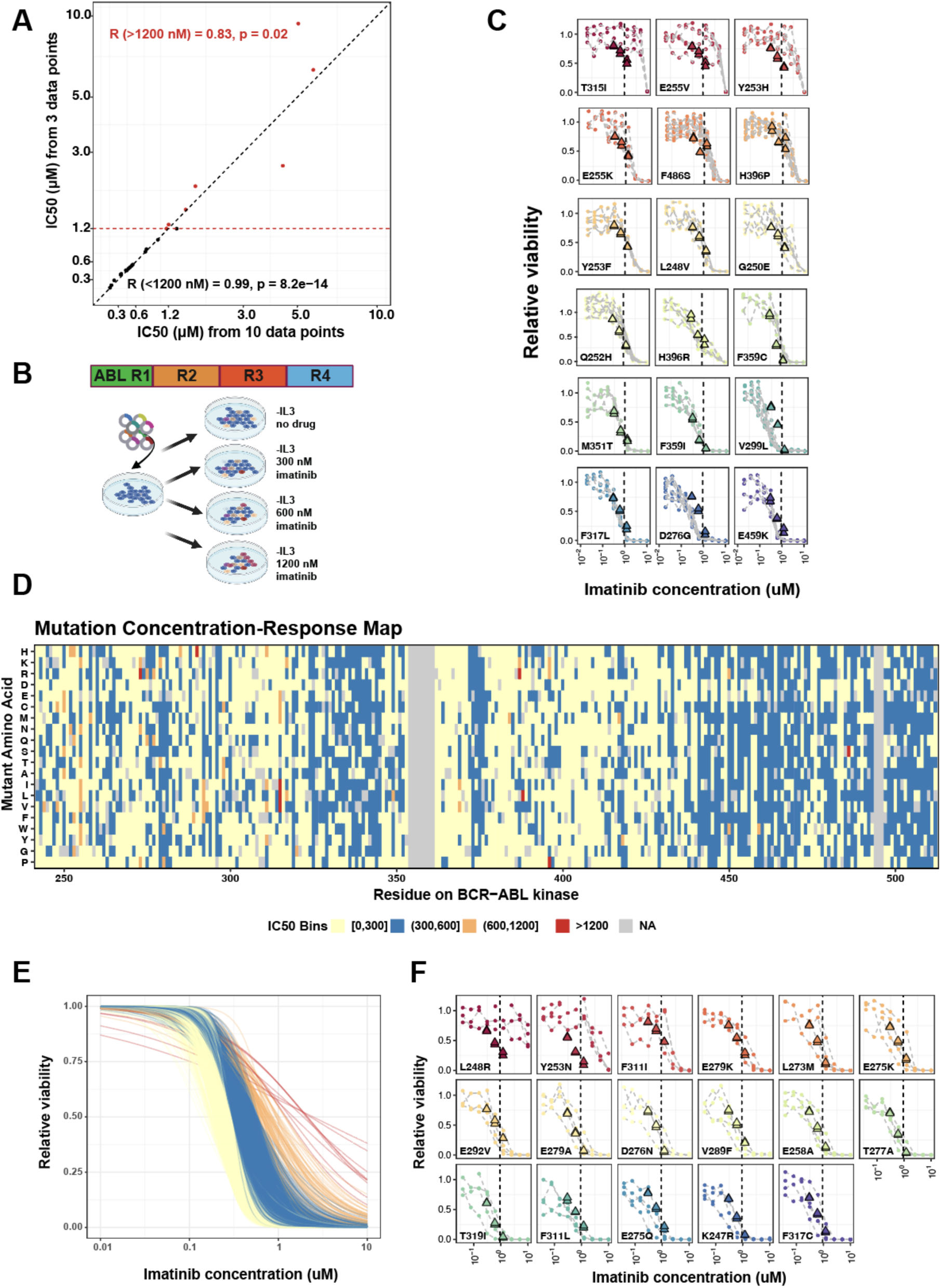
Imatinib Concentration Response Profiles for all BCR-ABL kinase variants validated using training and test mutant sets. (**A**) IC50 estimates from 3-point versus 10-point concentration response curves for n=21 mutant standards. (**B**) Deep Mutational Scanning of BCR-ABL kinase under no drug, and under 300nM, 600nM, and 1200nM imatinib. (**C**) Inferred mutant relative viabilities from DMS data (triangles, n=2 replicates) versus relative viabilities measured in single-mutant concentration-response curves (points, n=4 to 6 replicates) for 18 imatinib mutant standards. (**D**) DMS-inferred IC50 values for 4,923 BCR-ABL variants. (E) DMS-inferred concentration-response curves for 4,922 BCR-ABL variants (after excluding M351F due to poor model fit). (**F**): Inferred mutant relative viabilities from DMS data (triangles, n=2 replicates) versus relative viabilities measured in single-mutant concentration-response curves (points, n=4-8 replicates) for 17 moderate-to high-resistance mutants.

Building on this, we conducted a DMS screen of 4,923 single amino acid substitutions at three imatinib concentrations (300, 600, and 1,200 nM, Supplementary Tables S2-4) and a DMSO control (Fig. 3B, Supplementary Figures 4-5, Supplementary Table S5). We fit two-parameter logistic (2PL) models to generate concentration-response curves for 4,992 variants, excluding only one variant (M351F) due to a poor model fit (Fig. 3D). The framework was calibrated against a training set of 18 standards (Fig. 3C). For 14/18 variants within the target IC50 range (300-1,200 nM), we observed strong concordance between single-mutant and DMS-pool-derived viabilities. As expected, for the remaining four variants outside this range (with absolute IC50s >1200 nM or < 300 nM) we observed qualitatively consistent but less precise measurements. Then we externally validated against a test set of 17 new mutants, where we observed strong agreement for 15/17 variants (Fig. 3F). This process produced a high-confidence resource of concentration-response curves for ∼95% of all possible substitutions in the *BCR-ABL* kinase domain (Fig. 3E, Supplementary Fig. 6 shows concentration-response curves for our 18 gold standards).

### A clinical PK/in vitro PD Framework to Link In Vitro Data with Clinical Dosing

A major barrier in clinical oncology is the lack of a framework that connects in vitro drug response data (in vitro pharmacodynamics, PD) to patient-specific drug exposures (clinical pharmacokinetics, PK). To bridge this gap, we aimed to develop a PK/PD model that quantitatively links our high-throughput concentration-response data to clinically relevant imatinib exposures in patients. We used three key PK thresholds based on published steady-state concentrations: 444 nM for the standard frontline dose of 400 mg once daily (QD), 760 nM for an escalated dose of 400 mg twice daily (BID), and 916 nM for 500 mg BID⁶² (Fig. 4A).

**Figure 4.**
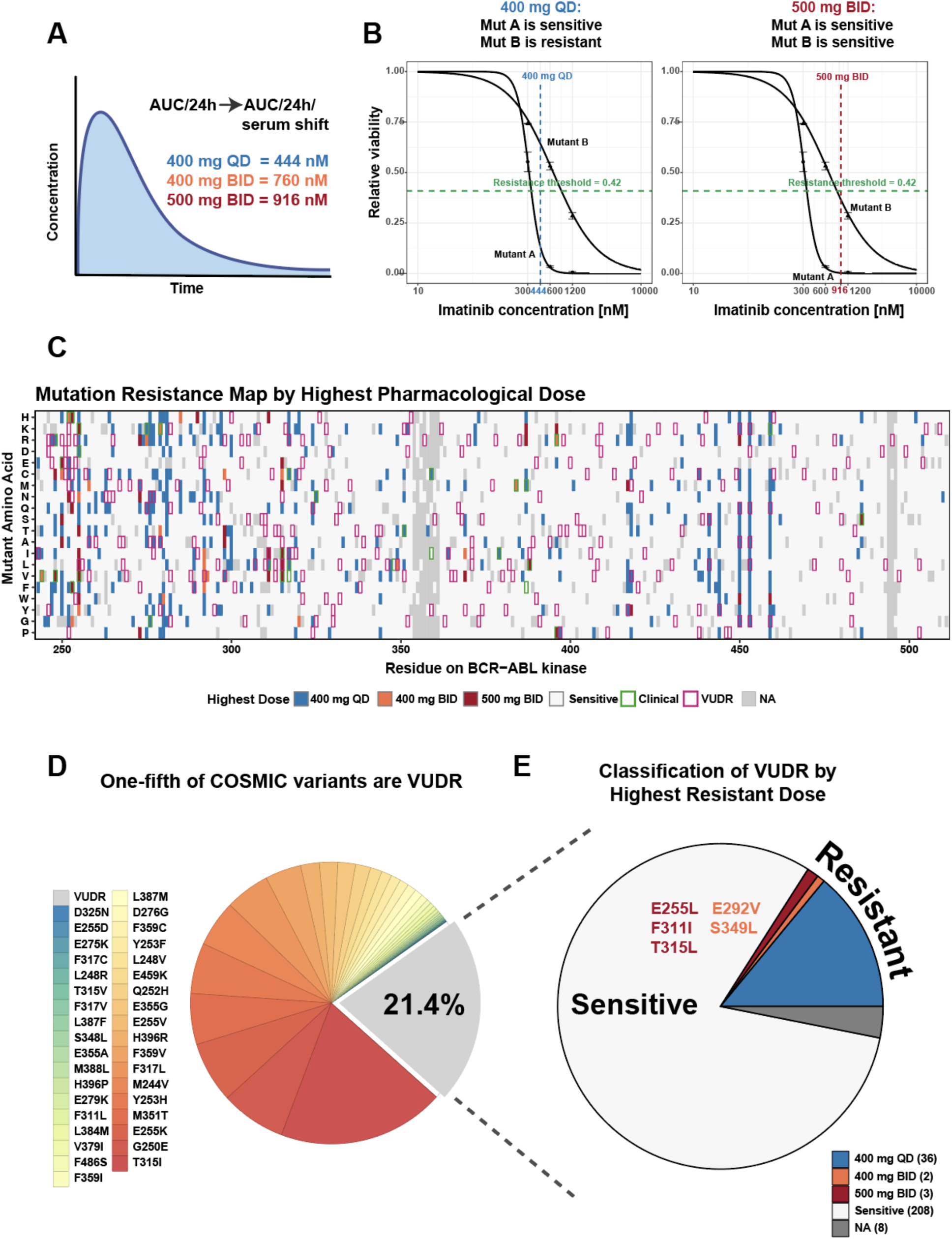
Linking patient-specific drug exposures (clinical pharmacokinetics, PK) to in vitro variant-specific resistance (in vitro pharmacodynamics, PD) enables dose-level classification and clinical interpretation. (**A**) In vitro drug responses were mapped to clinically relevant pharmacokinetic parameters, enabling estimation of serum-adjusted drug exposure at therapeutic doses (400 mg QD = 444 nM, 400 mg BID = 760 nM, 500 mg = 916 nM). (**B**) Relative viabilities were used to classify each of the 4,923 mutations as resistant or sensitive at clinically relevant doses, using optimal classification threshold (resistance threshold) via the Youden index. Shown are two examples: Mutant A (classified as sensitive at both 400 mg once-daily (QD) and 400 mg twice-daily (BID)) and Mutant B (classified as resistant at 400 mg QD and sensitive at 400 mg BID). Confidence intervals at 300, 600, and 1200 nM indicate replicate variability in the observed net growth-derived viability measurements. (**C**) Heatmap depicting resistance of individual mutants by highest pharmacological dose. Mutants are colored by the highest imatinib dose at which they are predicted to be resistant: sensitive at all doses (white), resistant at 400 mg QD (blue), resistant at 400 mg BID (orange), or resistant at 500 mg BID imatinib (maroon). Known clinically resistant mutations are highlighted in green, while COSMIC Variants of Uncertain Drug Resistance (VUDR) are highlighted in magenta. (**D**) Sanger clinical sequencing data (COSMIC database) of treatment refractory CML patients (n=1,859 patients with mutations, n=293 mutants). Majority of COSMIC CML mutants have known resistance mutants, and one-fifth of mutations are Variants of Uncertain Drug Resistance (VUDRs). Variants are ordered and colored by observed clinical abundance, with brighter red indicating higher frequency. (**E**) Most VUDRs are sensitive to imatinib, while ∼14% confer resistance to frontline imatinib dose (400 mg QD). Several VUDRs are resistant at higher doses, with two resistant at 400 mg BID and three at 500 mg BID.

We first developed a classifier to distinguish resistant from sensitive variants at the frontline 400 mg QD (444 nM) exposure. A logistic regression model was trained on a gold-standard set of 18 *BCR-ABL* mutations with validated clinical resistance status, including additional four sensitive mutations (total of 22). Using each variant’s relative viability at 444 nM, a logistic regression was trained to predict clinical resistance versus sensitivity, and an optimal classification threshold was defined using the Youden index (Fig. 4B; Supplementary Fig. 7). The resulting classifier performed with high accuracy, correctly classifying 20 of 22 mutations (V299L and F359I were misclassified) and achieving 94.1% sensitivity and 80.0% specificity (Supplementary Table S7; Supplementary Fig. 8). To ensure generalizability and guard against overfitting, we validated the classifier on an independent test set of 15 resistant mutations from patient data that were not used in training. The model correctly classified 13 of 15 variants, misclassifying only F317V and L387F as sensitive (Supplementary Table S9). This confirmed the classifier’s robust predictive performance.

With a validated classifier, we generated a global resistance map by applying it to all 4,922 variants at each of the three PK-defined thresholds (444 nM (400mg QD), 760 nM (400mg BID), and 916 nM (500mg BID)). This approach allowed us to model the clinical effect of dose escalation, identifying variants that could be re-sensitized at higher drug concentrations (Fig. 4B, right panel). The resulting map highlights resistance clustering at known clinical hotspots (e.g., G250, E255, T315) and provides a comprehensive annotation for each variant (Fig. 4C). Globally, 4,475 variants (∼91%) were sensitive at all tested exposures. The remaining 447 variants conferred measurable resistance at clinical doses: 382 were resistant at the 400 mg QD dose, 24 at 400 mg BID, and 41 at 500 mg BID. Of these resistant variants, 89 (∼23%) can arise from single-nucleotide variants (SNVs), making them more likely to be observed in patients.

The primary application of our framework is the functional annotation of Variants of Uncertain Drug Resistance (VUDRs). We identified 257 VUDRs from patient sequencing databases, which represent ∼21% of all clinically observed mutations in the dataset (Fig. 4D, Supplementary Table S8). Our analysis provided a definitive classification for these variants. The vast majority (∼81%) were sensitive to imatinib at the frontline dose, suggesting they are functionally inert passenger mutations. However, about 14% conferred resistance at standard dosing. A small subset of these remained resistant even at escalated doses, including 2 at 400 mg BID (E292V, S349L) and 3 at 500 mg BID (E255L, F311I, T315L) (Fig. 4E; Supplementary Table S10). This systematic classification enables confident interpretation of VUDRs and identifies cases where inexpensive imatinib dose escalation may be a viable clinical strategy over switching to an alternative TKI.

To facilitate the translation of these findings, we have made the entire resistance map publicly available as an interactive tool at: https://amplicomics.shinyapps.io/bcr_abl_resistance_map_by_clinical_dose/.

### Germline Variation Creates Ancestry-Specific Pathways to Resistance

To investigate how inherited genetic variation might alter pathways to drug resistance, we analyzed population-level data from the gnomAD database^61^, where approximately 18% of individuals carry at least one germline single nucleotide polymorphism (SNP) in the *ABL1* gene (Fig. 5A-C). Our analysis focused on identifying cases where a germline SNP converts what would be a low-likelihood multi-nucleotide variant (MNV) from the reference genome into a high-likelihood single-nucleotide variant (SNV) that confers resistance. A key example is the synonymous SNP T315T (rs1338400730), common in European populations, which enables the somatic acquisition of the highly resistant T315M mutation via a single nucleotide change -a pathway not accessible from the reference sequence (Fig. 5D).

**Figure 5.**
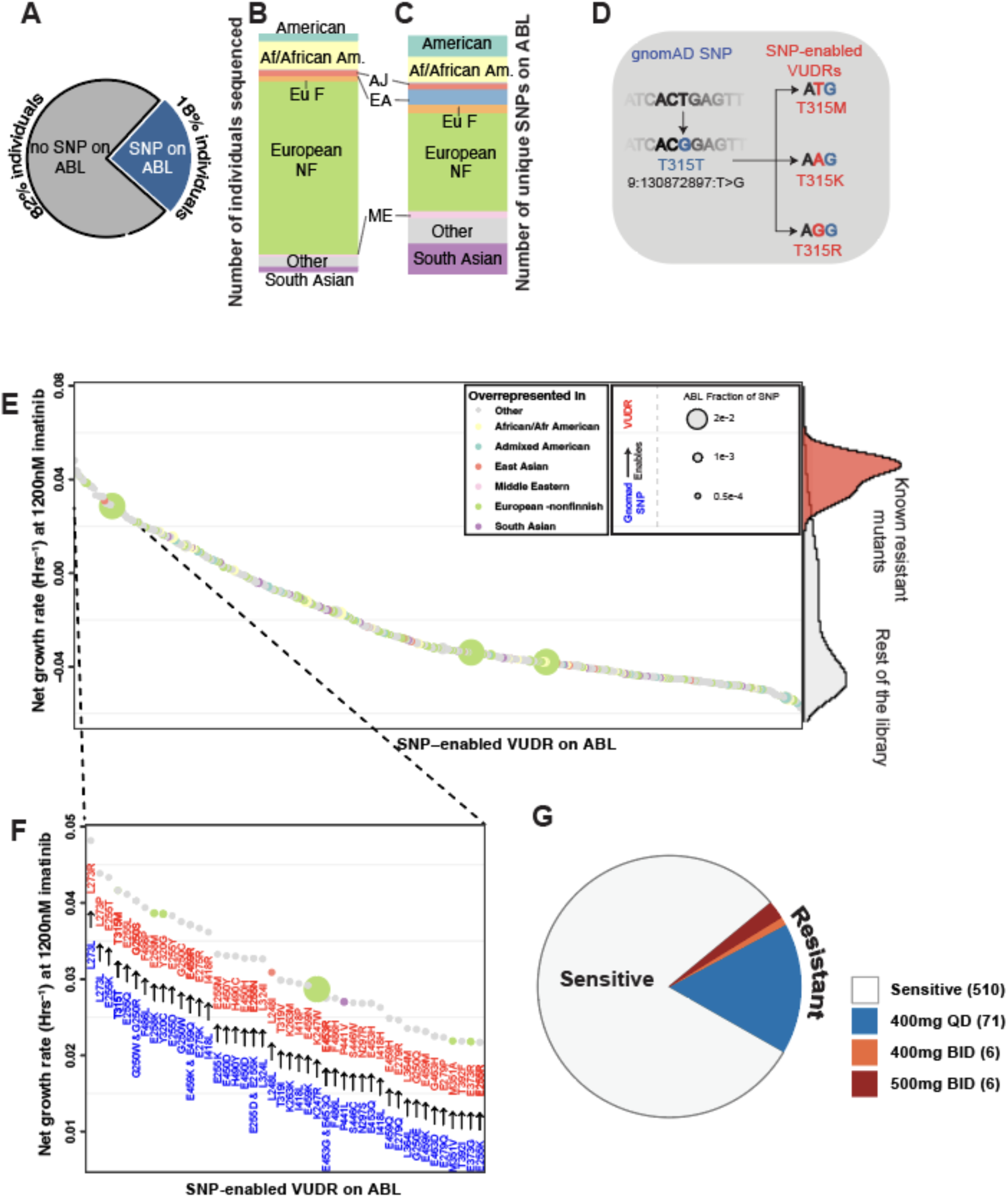
Germline variation suggests clinically actionable rare resistance mutations in specific genetic backgrounds. **(A)** (left) Fraction of gnomAD individuals with SNP in ABL (n=799,705). (**B,C**) These SNPs were detected across individuals from various ancestries (ME: Middle Eastern, Eu NF: European non-Finnish, Eu F: European Finnish, AJ: Ashkenazi Jewish, EA: East Asian,). **(D).** Schematic showing that population SNPs on ABL (blue), can convert low-likelihood MNVs into higher-likelihood SNVs (red). (**E)** Gnomad-enabled VUDRs (n=597 mutants) have a variety of imatinib resistance phenotypes as measured by growth rate at 1200nM imatinib. (**F**) The top 50 gnomAD-enabled VUDRs measured by resistance at 1200nM imatinib (**G)** Resistance characterization of gnomAD enabled VUDRs at clinically relevant imatinib doses.

Expanding this analysis across the *ABL1* coding sequence, we identified 182 gnomAD SNPs that create single-step evolutionary pathways to 596 otherwise inaccessible Variants of Uncertain Drug Resistance (VUDRs). By applying our resistance classifier, we determined that 83 of these 596 “germline-enabled” VUDRs are clinically resistant at therapeutic imatinib doses (Fig. 5E-G). Of these 83 resistant mutants, 71 confer resistance only at the approved 400mg daily dose and may be overcome by dose escalation.

These enabled resistance pathways can be ancestry-specific. For example, the E373D and K247W SNPs, which are overrepresented in European populations and enable the resistant E373H and K247R mutations. Similarly, the R460C SNP, overrepresented in African/African American populations, enables the resistant R460Y mutation. Notably, of the 12 VUDRs that remained resistant at doses above 400 mg once daily, 7 are located at known imatinib resistance hotspots (residues 250, 255, and 315), underscoring the clinical relevance of these germline-somatic interactions.

## DISCUSSION

The functional evaluation of VUS in oncogenes has been an active area of research^19,62–69^. However, assessing the drug resistance of VUS is a more challenging problem than simply adding a drug to existing DMS workflows. We rename this problem Variants of Uncertain Drug Resistance (VUDRs) to clearly distinguish the unique challenge of translating DMS studies of VUS to drug responses in people at clinical doses. The five main technical challenges that are lacking in the literature for the accurate classification of drug resistance in VUDRs include: 1) the lack of known clinical standards^9,45^, 2) the lack of a demonstrated ability to recover quantitatively accurate data without using those standards (especially considering known issues with sequencing errors)^54,59^, 3) not correcting for well-known time dependent changes in log2 fold change values when the overall pool fitness changes during an experiment (Supplementary Figure 2), 4) lack of in vitro responses across multiple drug concentrations that can be quantitatively related to patient pharmacokinetics, and 5) not establishing PK-PD relationships that can quantitatively relate human exposures to in vitro responses.

To address these challenges, we developed a strategy that combines Deep Mutational Scanning at multiple concentrations with a TileSeq-based readout to functionally assess nearly 5,000 BCR-ABL kinase domain variants across 3 clinically relevant concentrations of imatinib. Through a careful analysis utilizing standards, we show that we can create PK/PD relationships using high throughput data. We present our recipe in the diagram in Supplementary Figure 9 that outlines the five-step VUDR workflow for classifying the clinical drug resistance of BCR-ABL protein variants. Briefly, in the first step, a comprehensive library of single amino acid substitutions is generated for the target protein (e.g., BCR-ABL). In the second step, the variant pool is exposed to multiple concentrations of a small molecule (e.g., imatinib). Mutational counts are measured using TileSeq, a sequencing method that enables accurate quantification of variant frequencies. Subsequently, careful measurement of net growth rates, by integrating mutant allele frequency (MAF) from sequencing with time-matched cell density, eliminates artifacts typically seen in pooled log2 fold change analyses. In the third step, key known clinical resistance mutations are tested individually in single-mutant assays to validate the pooled DMS data (these standards could also be assayed as a validation exercise after screening). If the panel of concentration-response curves confirms the reproducibility of the pooled measurements then we can proceed. In the fourth step, for each variant, a simple concentration response model is fit to the concentration-response data. The mutation concentration-response map summarizes IC50 estimates across the protein, providing a landscape of concentration-response for each variant across the protein. Finally, each variant’s resistance status is classified based on its inferred response at clinically relevant serum-adjusted doses of a drug (e.g.,imatinib 400 mg once daily, imatinib 400 mg twice daily, imatinib 500 mg twice daily). The final heatmap assigns resistance classification per variant, enabling a clinical dose-aware interpretation of variant drug response. After using this workflow, we achieved measurements for 4,922 variants (96% of the intended region). The majority of missing variants were localized to one small region (354-361 aa) that failed to appear in our cloned library.

The work presented in this paper aimed to translate VUDRs in the BCR-ABL kinase domain into potentially clinically actionable interpretations. Interestingly, after >20 years of BCR-ABL studies, only a small subset of the most prevalent missense mutations in BCR-ABL have been systematically characterized for drug responses in vitro, but 257 VUDRs have been observed in the COSMIC database and remain unclassified in terms of clinical relevance to imatinib (or other drug treatment). We have measurements for 249 (97%) of those variants. We find that only a small percentage of VUDRs (41/257, ∼16%) are functionally resistant to imatinib at 400 mg QD, 400 mg BID, or 500 mg BID, suggesting that many variants in the COSMIC database are passenger mutations that are unlikely to be causative. This low percentage of functional variants is consistent with previous MAVE and SGE studies in other oncogenes, where only a small fraction of VUS were classified as loss-of-function - 14.6% in *PTEN*^70^, 18% in *TP53*^70^, and 25% (64/254) in *BRCA1*^19^. This is an interesting concordance given the large differences in disease/gene/context and strongly implies caution interpreting rare variants in known disease genes without calibrated data on the functional significance of those variants.

In our study we chose Ba/F3 cells because of their longstanding history as an acceptable model for tyrosine kinase addiction and drug treatment^71–74^. We also previously validated the ability to establish PK/PD relationships with limited numbers of ABL/KIT/EGFR mutants in this exact cell line^75^. However, Ba/F3 cells are a transformed murine cell line that does not encompass the complexity of human disease. Despite the obvious differences in cell context, we stress that we have 94.1% sensitivity and 80% specificity in our study for predicting resistance variants within the kinase domain. We also validated small panels for accurate PK/PD relationships and prediction in a previous paper and achieved 90.3% sensitivity and 84.2% specificity^75^. However, these values for sensitivity and specificity relate only to variants in the kinase domain and variants elsewhere in the genome have unvalidated performance characteristics. Despite these caveats, prior work on many tyrosine kinases and many tyrosine kinase inhibitors supports the use of the Ba/F3 model as a simple model of oncogene addiction and inhibitor sensitivity that extends to many oncogenes^38,76–81^ and has surprising accuracy in discovering resistance variants within the oncogene.

Our work provides a strong clinical and financial rationale for re-evaluating the role of imatinib in the context of variant-driven resistance, given the high cost and varying safety profiles of available ABL tyrosine kinase inhibitors (TKIs)^82–86^. TKIs are expensive, and while many patients who progress on imatinib switch to another TKI, imatinib is currently generic. Thus, in addition to having the most clinical safety data and being well regarded for its safety profile relative to drugs like nilotinib and ponatinib, imatinib is also less financially toxic than branded TKIs. In fact, generic imatinib remains the most cost-effective TKI option, with annual costs as low as $200, while generic dasatinib ranges from $3,000 to $100,000, and patented second-to fourth-generation TKIs range from $20,000 to $300,000 annually, despite no clear survival advantage over imatinib^87^. Imatinib has been evaluated at higher doses and has proven safe and effective at these higher doses^88–93^. Our concentration response data suggests the unique possibility that many imatinib VUDR might be treated with dose escalation of this safe, cheap, and effective agent. Beyond what we have established with imatinib, there are now 6 approved ABL TKI. All have different resistance profiles for common resistance variants and different side effect profiles. Expanding data sets like this to all BCR-ABL TKIs in the future could allow for the principled selection of the correct TKI for the correct VUDR, and when multiple TKIs might address the same VUDR, it may allow for more flexibility in choosing the right TKI based on other clinical criteria.

Our results also highlight the importance of integrating population-level genetic diversity into DMS-based models of mutant fitness. In specific ancestry groups, these variants exist as accessible SNVs and may meaningfully impact treatment outcomes in patients when they lead to rare mutations that are not accessible in a standard genome. We were surprised by the presence of single step mutations capable of producing the highly resistant and somewhat rare T315M mutation in European Non-Finish population background, and the ability of germline variants such as R460C to create the unique resistance mutation R460Y specifically in the African American population. As generic drugs expand TKI accessibility, and Novartis continues to make drugs accessible in countries that lack a commercial supply, more people around the world with diverse germline sequences will develop TKI resistance. Our measurements may help in N of 1 patients in rare backgrounds to effectively navigate therapy in resource poor settings.

Ultimately, as the FDA allows labels to include rare variants for which only in vitro data exists, and the NCCN guidelines frequently incorporate diverse lines of evidence with diverse quality into treatment recommendations, this work takes a big step towards the possibility that the FDA and NCCN could incorporate high throughput data into labels or evidence blocks in order to help clinical decision making for patients with rare mutations. To this end, we developed a publicly available interactive Shiny app https://amplicomics.shinyapps.io/bcr_abl_resistance_map_by_clinical_dose/ as a mutation-guided support tool built on a high-throughput resistance mutation map to help disseminate our results to clinicians and researchers. Our goal is to expand this tool across all ABL TKIs to enable informed and personalized treatment selection, including potential combination therapies. Building on the paradigm shift observed in functional genomics for interpreting VUS, our work marks a similar transition toward systematically characterizing clinical drug resistance of VUDR in cancer.

## METHODS

### Plasmid design and mutagenized library preparation

A plasmid encoding the full-length BCR-ABL fusion protein (6,096 bp) was synthesized on the pUltra backbone (Addgene #24129) and sequence-verified (Genscript). The construct contains a 2A self-cleaving peptide to ensure balanced multicistronic expression.

Mutagenesis was performed by Twist Bioscience using the single-site variant library (SSVL) platform, generating all possible missense substitutions between residues 242-512 of the ABL kinase domain (270 residues; 5,119 variants). The library was distributed across a 96-well plate with eight residues mutagenized per well.

The library was subdivided into four regions (ABL 242-321, ABL 322-393, ABL 394-465, and ABL 466-512), which were transduced and processed separately. Each region was transformed into NEB Stable *E. coli*, yielding >1,000 colonies per variant (>5 × 10^6^ total colonies across fifty 15-cm plates). Colonies were grown at 30 °C for 48 h, and plasmid DNA was purified using E.Z.N.A. Plasmid Midi Kits (Omega Bio-Tek), yielding ∼3 mg per preparation.

### Transfection and infection of the mutagenized library

Because the full-length BCR-ABL construct was 6,096 bp, the transfer plasmid spanned ∼10.5 kb between the 5′ and 3′ LTRs, near the upper limit of lentiviral packaging. Transfections were therefore optimized for efficiency. The plasmid library (transfer vector) and four packaging plasmids were co-transfected into HEK293T cells using Lipofectamine 3000 (Thermo Fisher Scientific). For each 10-cm dish, 35 µg of transfer plasmid and 2.5 µg of each packaging plasmid were combined at a 5:1 Lipofectamine:DNA ratio. Medium was replaced 12 h after transfection, and viral supernatant was collected at 36 h and 72 h. Excess virus was concentrated using Lenti-X Concentrator (Takara Bio), resuspended in PBS, and stored at - 80°C.

Ba/F3 cells were infected with lentivirus in the presence of 6 µg/mL polybrene (Sigma-Aldrich). After 36 h, cells were washed and resuspended in fresh RPMI medium (Thermo Fisher Scientific). Cells were allowed to recover for 3 d prior to downstream selection. Infection efficiency was monitored to ensure a multiplicity of infection (MOI) < 1. To maintain library representation, ∼2 × 10^8^ Ba/F3 cells were infected, targeting >1,000 transduced cells per variant (>5 × 10^6^ cells total).

At 72 h post-infection, GFP-positive cells were enriched by fluorescence-activated cell sorting (FACS) to >95% purity. The sorted population was expanded until sufficient cells were available for imatinib selection. Baseline sequencing of the transduced Ba/F3 library, compared to the plasmid DNA input, showed minimal loss of library evenness (data not shown).

### Screening the library

At Day 0 (D0), 30 million cells were pelleted for each cell line and each replicate. Selection experiments were initiated such that 5 million cells were seeded at ∼10% confluence. Prior to screening, concentration-response assays were performed using Ba/F3 cells expressing BCR-ABL wild type (WT) and clinically validated resistant mutants to establish relevant drug concentrations. For Ba/F3 BCR-ABL cells, the library was selected at the IC25, IC50, and IC75 concentrations of imatinib (300, 600, and 1200 nM, respectively). Selection was initiated by simultaneous withdrawal of IL-3 and addition of imatinib.

Cells were maintained in exponential growth by resuspension into fresh drug-containing media every 2-3 days and splitting to maintain <50% confluence (∼1 × 10^6^ cells/mL). Daily measurements of viability, cell density, and mCherry positivity were performed using a BD Accuri C6 Plus flow cytometer. For each timepoint, cell density and dilution factors were recorded and later incorporated into growth rate calculations.

Selection experiments were conducted for 6 days. At the final timepoint (Day 6), ≥30 million cells were pelleted for each condition. When viability was <90% at Day 6, live cells were enriched using Ficoll-Paque (Cytiva). As a positive control for resistance, T315I-2A-mCherry cells were spiked into the library at a frequency of 1:2000.

### Duplex sequencing library preparation and analysis

Our scalable duplex sequencing workflow uses a combination of the original duplex sequencing protocol^94^ and the CRISPR-DS protocol^95^. The overall workflow includes extracting genomic DNA (∼1 hour prep time), generation of gRNA duplexes (∼1 hour prep time), targeted Cas9 cleavage of genomic DNA (∼1 hour digest), enrichment of low molecular weight DNA with spri (∼1 hour bead cleanup), end-repair and A-tailing of DNA (∼1 hour prep time), ligation of duplex sequencing adapters (∼1 hour ligation step), PCR1 to index and amplify libraries (∼2 hour PCR), hybridization with biotinylated ABL oligos and blockers^96^ (overnight incubation), targeted enrichment of ABL using streptavidin beads (∼3 hours), on-bead PCR2 to amplify final libraries (∼1 hour PCR), and cleanup of the duplex sequencing library with SPRI beads.

Each duplex sequencing library was sequenced on an Illumina NovaSeq platform (baseline and the DMSO arm of Day 6 libraries (Region 1) were previously published in Sokirniy, Inam et al., 2025^97^. The amount of sequencing required per library depends on the estimated complexity of the library. Generally, for a duplex sequencing library prep that uses 20ug of input DNA, 200 million 150bp paired ends reads should suffice. Whether or not enough sequencing was performed can be confirmed by looking at the count distribution of the duplex barcodes. The key metric is the median number of times each duplex barcode is seen post-sequencing, also known as the peak tag family size. A peak tag family size of 20 means that the library is sequenced sufficiently.

We used a custom bioinformatics pipeline to analyze the duplex sequencing data. Briefly, dunovo (53) was used for tag-clustering (consensus calling) of duplex barcodes from the raw sequencing data. Next, consensus-called, error-corrected read pairs were aligned to the ABL CDS (NM_005157.6) using the bwa-mem2 aligner. Next, filtering steps were performed on the aligned reads to remove contaminating reads from the mouse genome. This was done by allowing a maximum of 5 mismatches in the MDZ field of the BAM file. Next, a custom variant caller was used for variant calling and variant annotation. The output of this bioinformatics pipeline had counts for each mutant detected, and sequencing depths at each residue of ABL. All consensus calling and alignment steps were performed in Linux and all variant calling and downstream data processing and analyses steps were performed in RStudio.

### Tileseq Library preparation and analysis

For TileSeq sample and region-specific primer design, the BCR-ABL cDNA kinase domain was split into 9 overlapping 150-bp tiles (2-3 per each subregion: Region 1: ABL 242-321 (tile 1_1, 1_2, and 1_3), Region 2: ABL 322-393 (tile 2_1 and 2_2), Region 3: ABL 394-465 (tile 3_1 and 3_2), and Region 4: ABL 466-512 (tile 4_1 and 4_2). Each tile was amplified via PCR using 1 µg of baseline or post-treatment sample per reaction (30 cycles: 98 C for 15 s, 65 for 30 s, 72 C for 30 s) with two biological replicates per condition using long sample and tile-specific TileSeq primers (Supplementary Table S11). TileSeq libraries were pooled at equimolar concentrations and sequenced using paired-end 150 bp reads on the Illumina NovaSeq X Plus platform, generating over 1 × 10⁶ paired-end reads per sample (n = 88 libraries, Supplementary Table S12).

TileSeq sequencing libraries were processed by overlapping, error-correcting, and merging paired-end reads using PEAR v0.9.11^98^. Merged reads were aligned to the ABL coding sequence (NM_005157.6) using bwa-mem2^99^ in a Linux environment. Variant calling and annotation were performed with a custom R-based pipeline, generating counts and sequencing depth for each unique mutant at each ABL residue. Subregions and subsequent full-sequence BCR-ABL kinase for each condition, including the no-drug control, were reconstructed from tiled data using custom Python scripts based on tile position cutoffs.

### Predicting mutant-specific net growth rates from concentration response curves

Concentration response curves measure the fractional survival of a population as a function of the applied concentration. We first fit a two parameter logistic to the concentration response curves for each mutant:

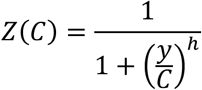

Where Ζ is the fractional survival parameter, y, is the drug concentration, and the parameters C and h are the IC50 and the hill coefficient, respectively.

Next, a drug kill rate, α, was derived using a formula previously derived by Zhao et al in 2016^100^:

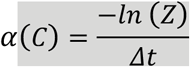

Where Δt is the treatment duration for the concentration response study. This drug kill rate is the concentration-dependent effect that the drug has on reducing the net growth rate of each mutant. For a given concentration, C, the drug kill rate and the growth rate of the cell-line without drug can be used to calculate the net growth rate in the presence of drug:

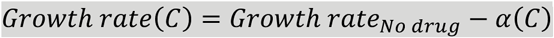

Exponentially growing BaF3 cells without drug have a growth rate of 0.055Hrs^-1^, which is a doubling time of 12-13 hours.

### Calculating net growth rates from sequencing data

Log fold-change (LFC), which is a measure of mutant enrichment/depletion, was calculated as the log₂ ratio of MAF at the final timepoint relative to the initial timepoint:

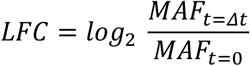

To estimate net growth rates of mutant populations, we inferred mutant counts at each timepoint. These counts were determined from MAFs and integrated with two key screen metadata metrics: cell density and dilution factor (DF) at the respective timepoint.

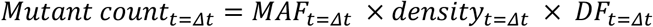

The cell density represents the total population density of the mutant pool at a given time, while the dilution factor accounts for the cumulative split ratio of the culture over the experiment duration. For example, at 𝑡 = 𝛥𝑡, if T315I was sequenced at a mutant allele fraction of 10^-4^, and was present in a flask at a cell density of 2 million cells/mL, and this flask was split at a dilution factor of 1:10 at some point between the start of the screen and 𝑡 = 𝛥𝑡, T315I would have an estimated 𝑀𝑢𝑡𝑎𝑛𝑡 𝑐𝑜𝑢𝑛𝑡*_t=Δt_* of 2,000 cells. Using these values, mutant growth rates were calculated as the natural logarithm of the ratio of mutant counts between two timepoints, normalized by the elapsed time interval (Δt):

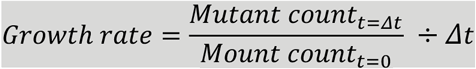

### Correction of DMS-derived net growth rates using linear regression

To align net growth rate measurements from deep mutational scanning (DMS) with those obtained from individual mutant assays, we applied a linear regression-based correction. Specifically, for each imatinib concentration (300 nM, 600 nM, and 1200 nM), we used a set of our experimentally validated 18 mutant standards to fit a linear model capturing the relationship between pooled DMS and single-mutant net growth rates. We verified that this model reduced the RMSE of our mutant standards (Figure 2C), and was subsequently used to transform the pooled DMS measurements (Supplemental Table S2), effectively correcting for systematic assay-specific deviations.

### Converting DMS growth rates to concentration-response curves

Net growth rates for each BCR-ABL variant were quantified from deep mutational scanning (DMS) experiments performed at 300 nM, 600 nM, and 1200 nM imatinib, and normalized to the wild-type net growth rate of BaF3 cells (0.055 Hrs^-1^) within matched drug-treated conditions to account for baseline growth differences. Variant-specific concentration-response curves were fit using dr4pl R package with four-parameter logistic (4PL) regression model (the minimum and maximum relative viabilities in this model were fixed to 0 and 1), enabling estimation of IC50 values, Hill coefficient (slope), model convergence status, and residual sum of squares (RSS) for curve fit quality.:

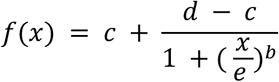

f(x) = fit (concentration-dependent response)

b = Hill slope

c = lower asymptote (minimum response)

d = upper asymptote (maximum response)

e = IC50

### Clinical resistance classification of BCR-ABL mutations using DMS concentration-response predictions

A set of 22 clinically annotated gold-standard resistant and sensitive mutants (including 18 mutants experimentally validated in the lab) was used to train logistic regression models to distinguish resistance status based on relative viability (Supplemental Figure 7). Classifier performance was evaluated on the gold-standard set, achieving 93% overall accuracy with 94.1% sensitivity and 80% specificity (Supplementary Table S7, Supplementary Fig. 7).

The optimal probability cutoff was determined using the Youden index and back-calculated to a relative viability value, defining the resistance threshold at the intersection of the logistic curve with the cutoff line (0.42):

At each dose, we computed the Youden index *J(t)* across candidate probability cutoffs *t* and took the maximizing *t* as the optimal threshold:

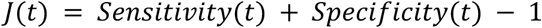

where:

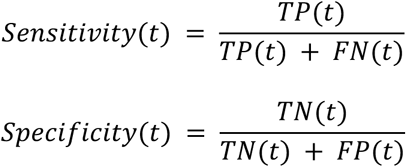

The optimal probability threshold was *t\**:

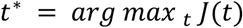

Predicted probabilities were obtained from a logistic regression of the binary resistance label Y ∈ {0,1} on relative viability x:

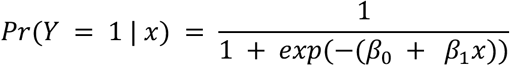

Let t* be the optimal probability cutoff; the corresponding threshold on relative viability (resistance threshold), x_cut_, is the intersection of the fitted logistic curve with the horizontal line p = t*:

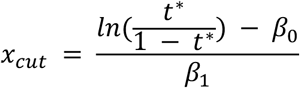

Variants were classified as resistant if *p_i_ > t\** and sensitive otherwise.

To enable classification of all screened variants at clinically relevant drug exposures, this fixed resistance threshold (0.420968716) was applied to DMS measurements at serum-free equivalent concentrations corresponding to common clinical dosing regimens. We used recently estimated steady-state, unbound plasma concentrations for imatinib dosing schedules: 400 mg once daily (QD; 444 nM), 400 mg twice daily (BID; 760 nM), and 500 mg BID (916 nM)^75^. This approach enabled mapping of in vitro resistance profiles to therapeutic exposure levels.

### GnomAD analysis

We used the genome aggregation database (gnomAD v4.1.0, https://gnomad.broadinstitute.org/) to obtain population-level single nucleotide polymorphism (SNP) data for the ABL gene (n = 799,705 individuals). From this dataset, we identified 3,356 SNPs within ABL, of which 153 caused amino acid changes (missense mutations) specifically within the kinase domain. We excluded any variants that were frameshifts, insertions/deletions, or located outside of the coding sequence of the ABL kinase domain.

We then focused on multi-nucleotide missense variants (MNVs) in ABL that cannot be formed by a single nucleotide change relative to the reference ABL transcript (ENST00000318560). For these MNVs, we checked whether any of the missense SNPs observed in gnomAD could *enable* them - meaning that the gnomAD variant changes the original codon into one that could produce the same MNV with just one additional nucleotide change. This analysis identified 596 such “gnomAD-enabled” amino acid substitutions (Supplementary Table S13).

In other words, the 153 missense SNPs observed in gnomAD make it possible for 596 multi-nucleotide amino acid substitutions to arise via just one additional point mutation. These mutations and their resistance characterization is summarized in Supplementary Table S13.

## Data availability

Sequencing data from pooled DMS screens are available at Sequence Read Archive under BioProject ID PRJNA1279649.

## Code availability

The code for processing pooled DMS data and generating figures is available on GitHub at: https://github.com/pritchardlabatpsu/abl_dms.

## SUPPLEMENTARY FIGURES

**Supplementary Figure 1.**
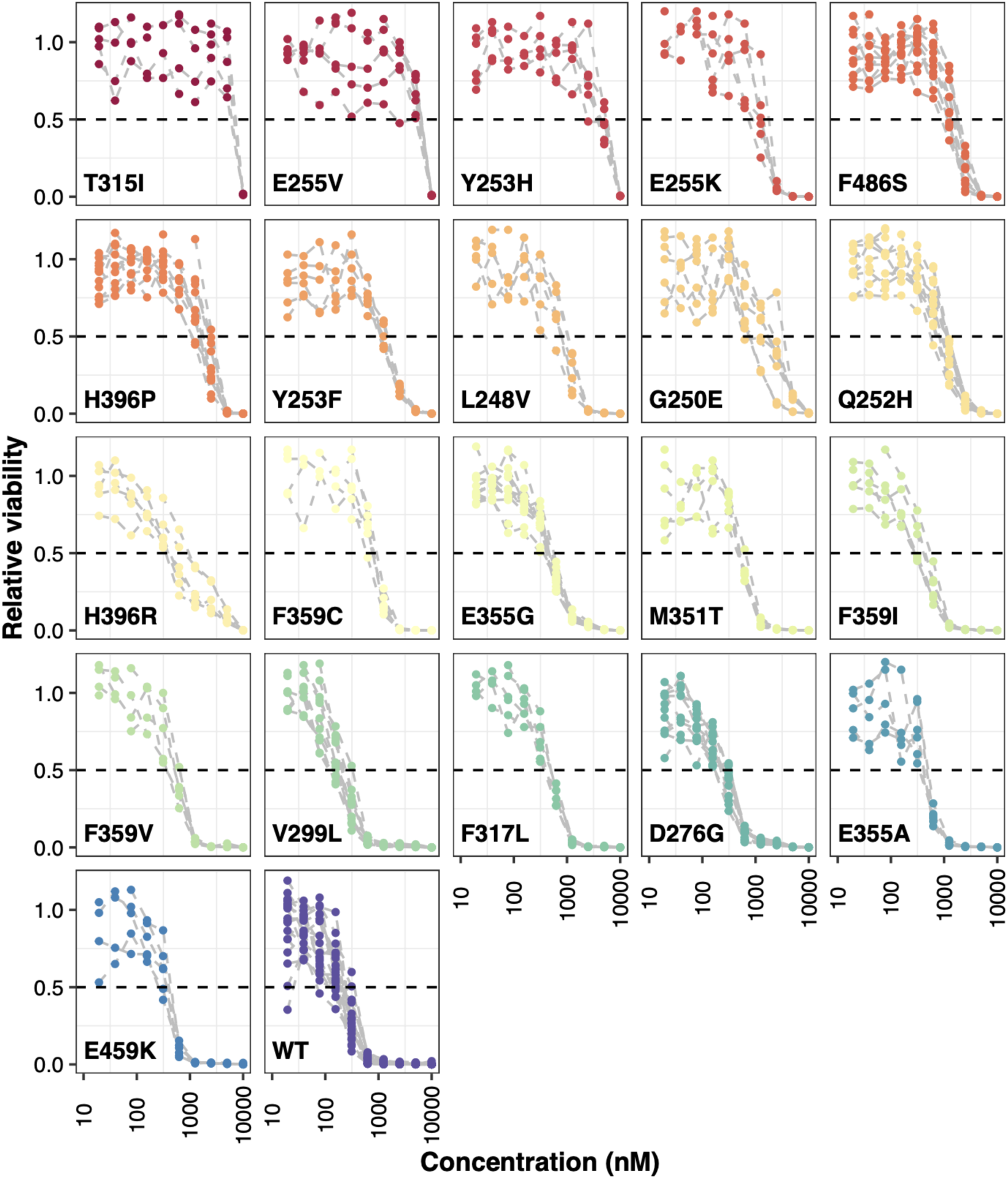
Concentration-response curves for 21 BCR-ABL mutants, including wild type (WT), tested across 10 concentrations of imatinib (n=4-8 replicates).

**Supplementary Figure 2.**
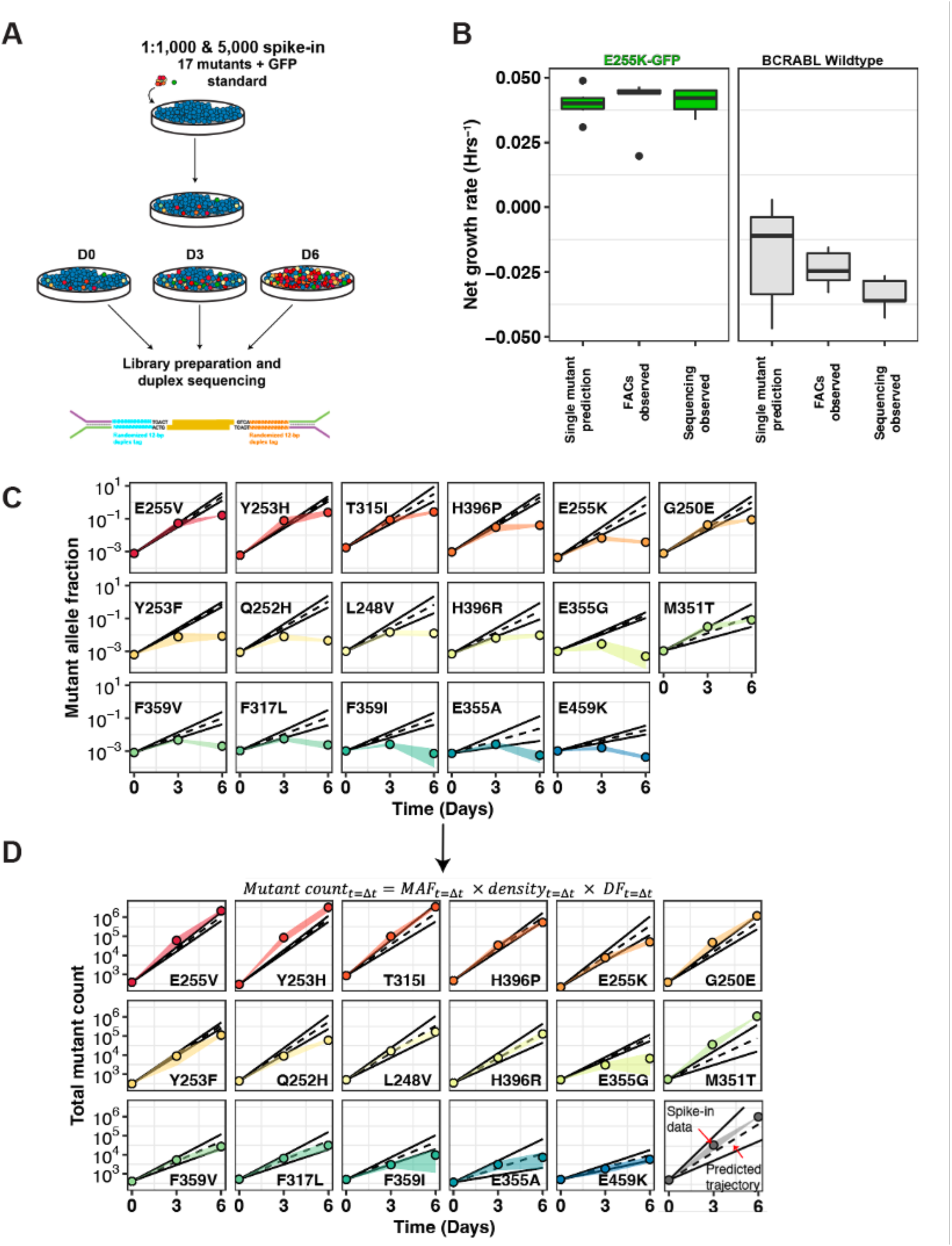
A: 17 imatinib resistant mutants in BCRABL were spiked-in at low depths of 1:3,000 to 1:5,000, screened and measured with duplex sequencing, **B:** Net growth measurements of BCRABL_E255K_-GFP and BCRABL_WT_ in a DMS pool versus single-mutant assay, measured by FACs and duplex sequencing, **C:** Measuring resistance by log2fold change for each of 17 imatinib resistant mutants, **D:** Trajectory of the 17 imatinib resistant mutants measured in a DMS pool with a correction method (colored points) versus dose-response predictions (black lines). The correction strategy to calculate net growth rates integrates mutant allele frequency (MAF) from sequencing with time-matched cell density measurements (density t = delta t). This correction removes log2fold change-driven distortions. The colored shaded region represents the standard deviation of duplex sequencing measurements across two screen replicates.

**Supplementary Figure 3.**
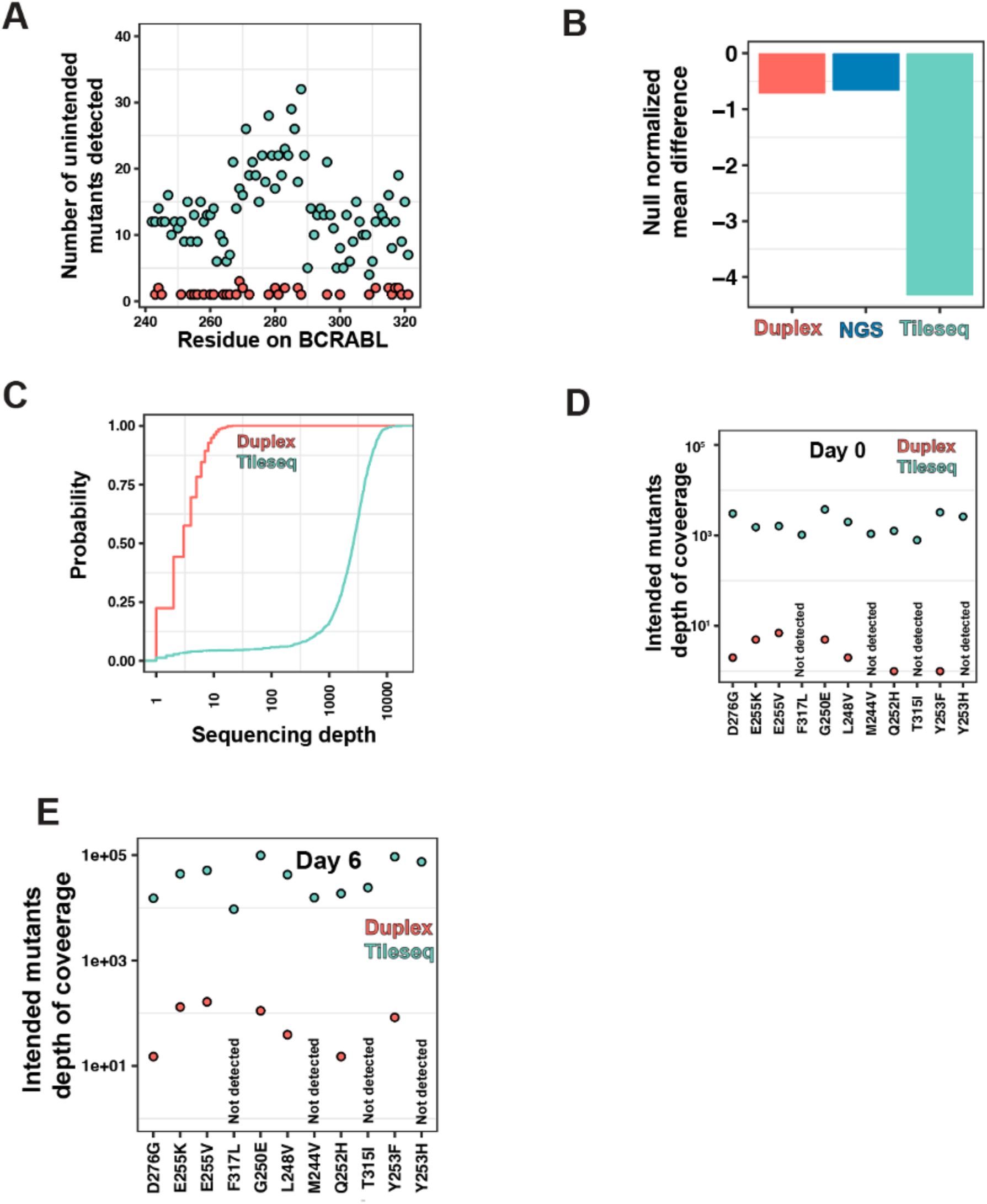
A: Number of unique unintended mutants detected by Duplex Sequencing (pink) and TileSeq (cyan) in region 1 of the BCRABL kinase. **B:** Null Normalized Mean Difference as a statistics to assess the separation of intended versus unintended codons using NGS, Duplex sequencing and TileSeq. The lower the value, the better the separation. C: Distribution of sequencing depths using Duplex Sequencing and TileSeq. D: Distribution of coverages of mutant standards, E: Distribution of the numbers of unique intended mutants detected by Duplex Sequencing and TileSeq.

**Supplementary Figure 4.**
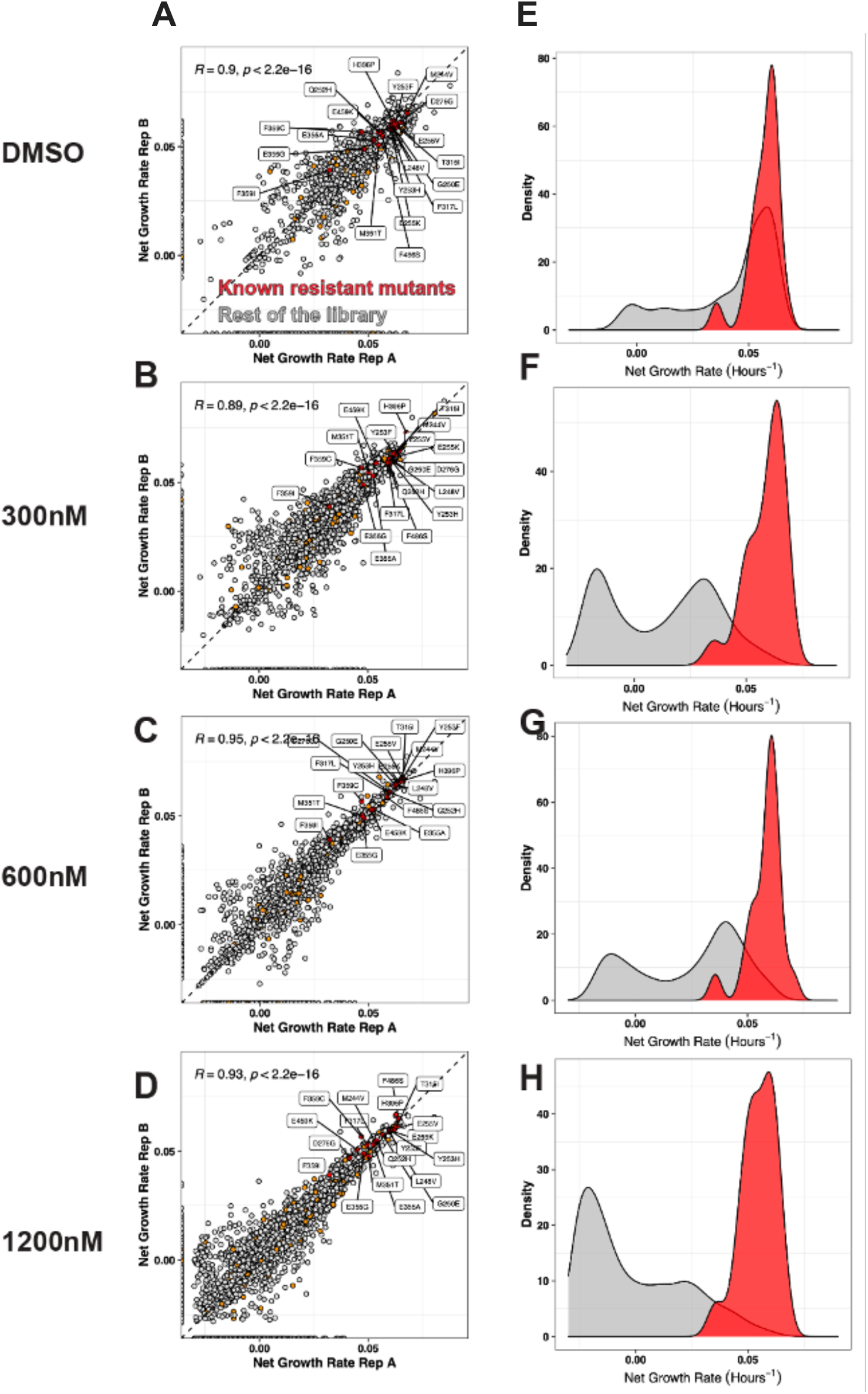
Biological replicate correlation of net growth rates and density plots of resistant mutants versus the rest from pooled DMS screen of BCR-ABL mutants under A, E: no drug, B, F: 300 nM imatinib, C, G: 600 nM imatinib, D, H: 1200 nM imatinib.

**Supplementary Figure 5.**
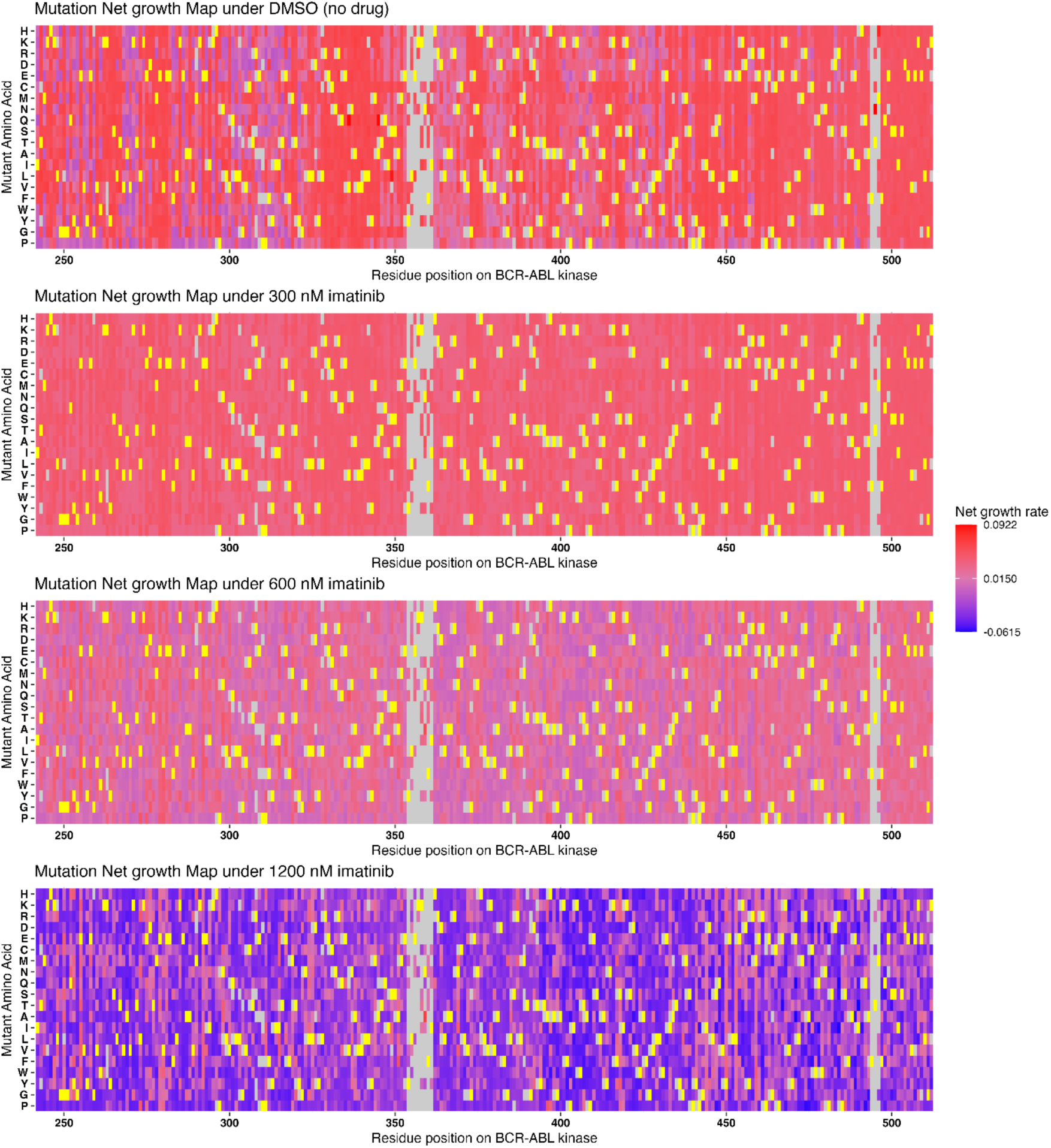
Net growth rate heatmap for 6-day BCRABL DMS screen under: A: no drug, B: 300nM of imatinib treatment, C: 600nM of imatinib treatment, and D: 1200nM of imatinib treatment. Yellow residues indicate WT amino acids.

**Supplementary Figure 6.**
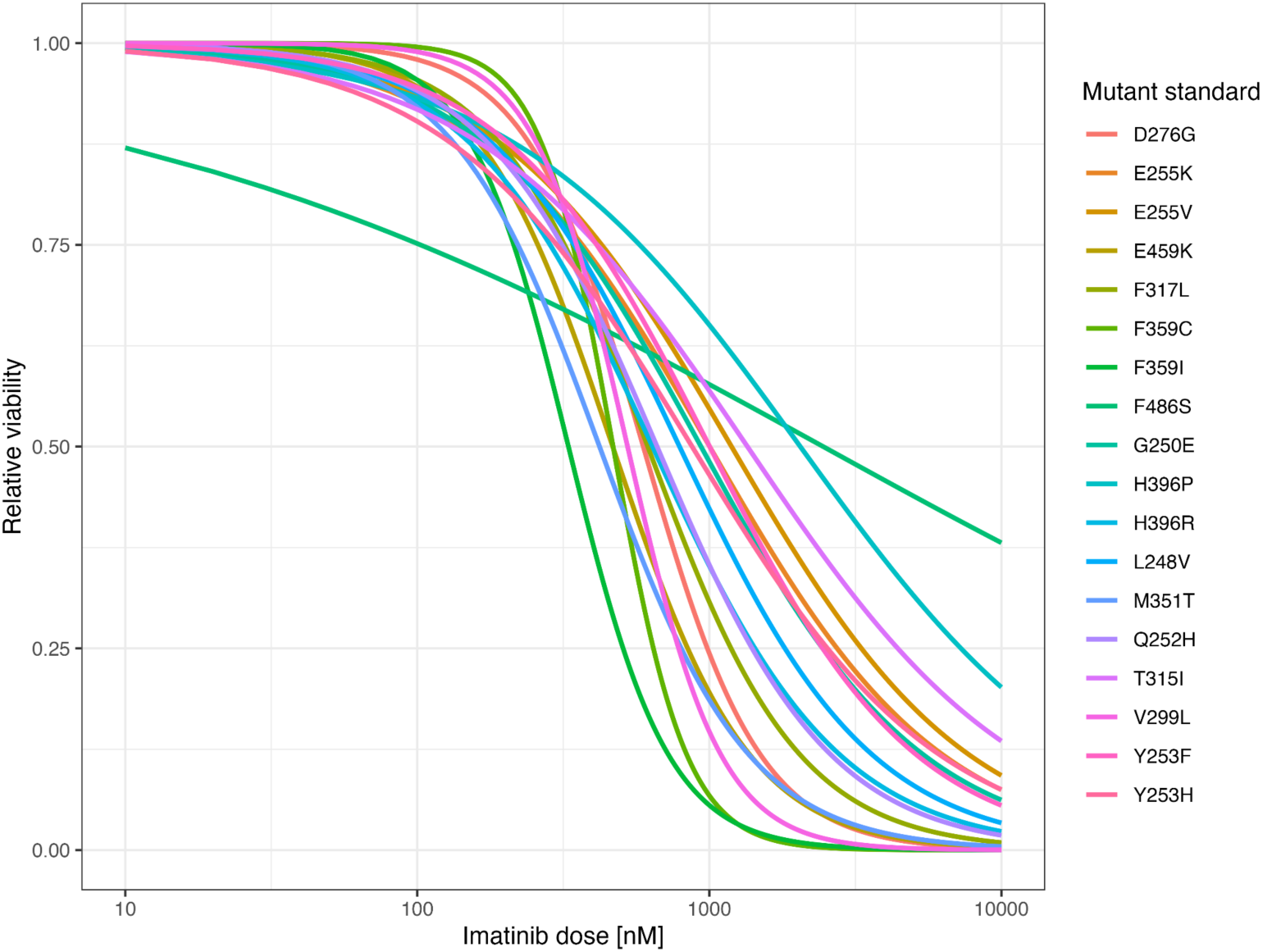
Concentration-response curves for 18 BCR-ABL mutant standards, tested across a range of imatinib concentrations (10-10,000 nM).

**Supplementary Figure 7.**
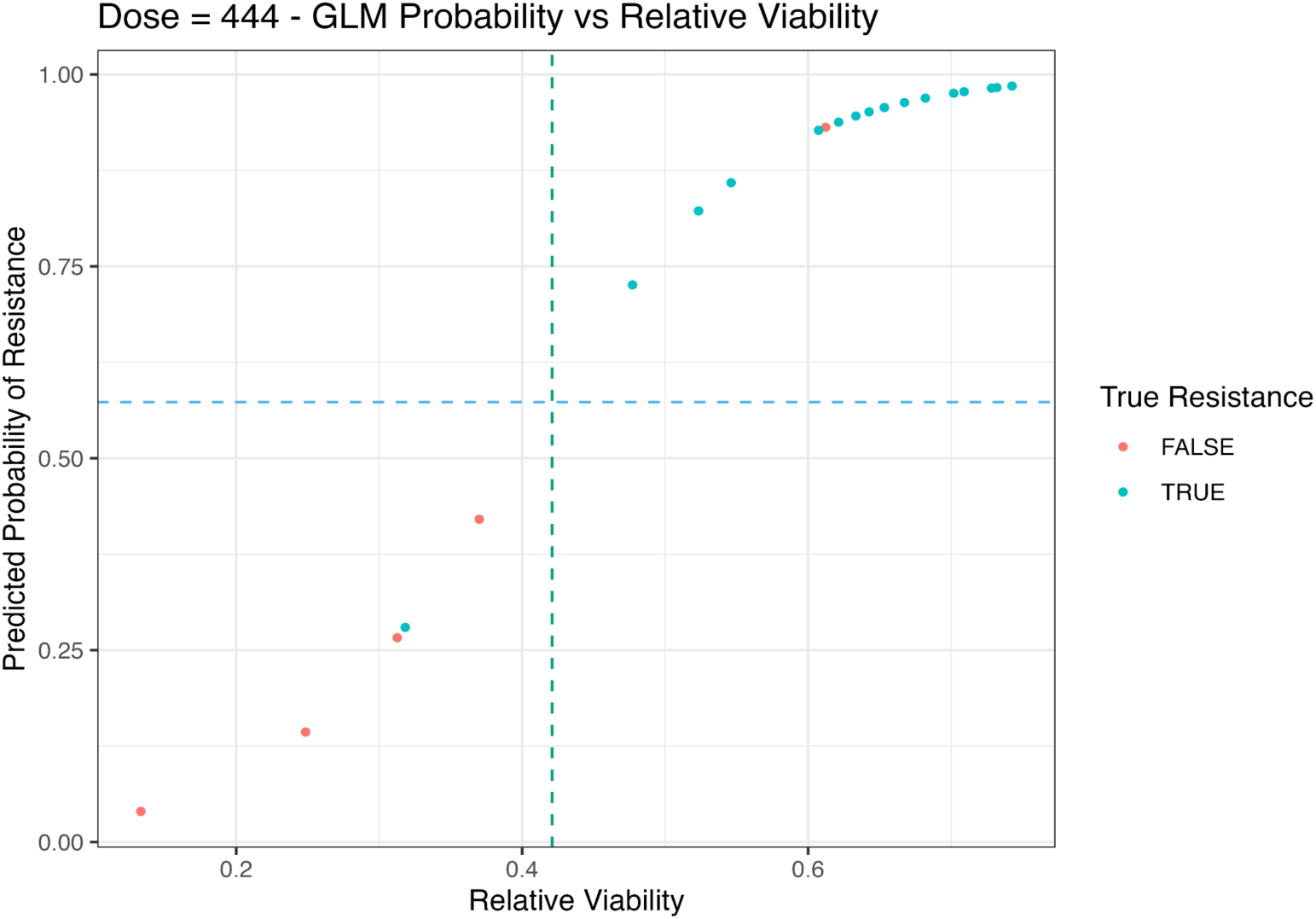
Logistic regression-based resistance threshold determination at the frontline imatinib dose. Logistic regression was used to model the probability of resistance as a function of relative viability. The optimal probability cutoff was identified using the Youden index and back-calculated to a relative viability value, defining the resistance threshold at the intersection of the logistic curve with the optimal probability line.

**Supplementary Figure 8.**
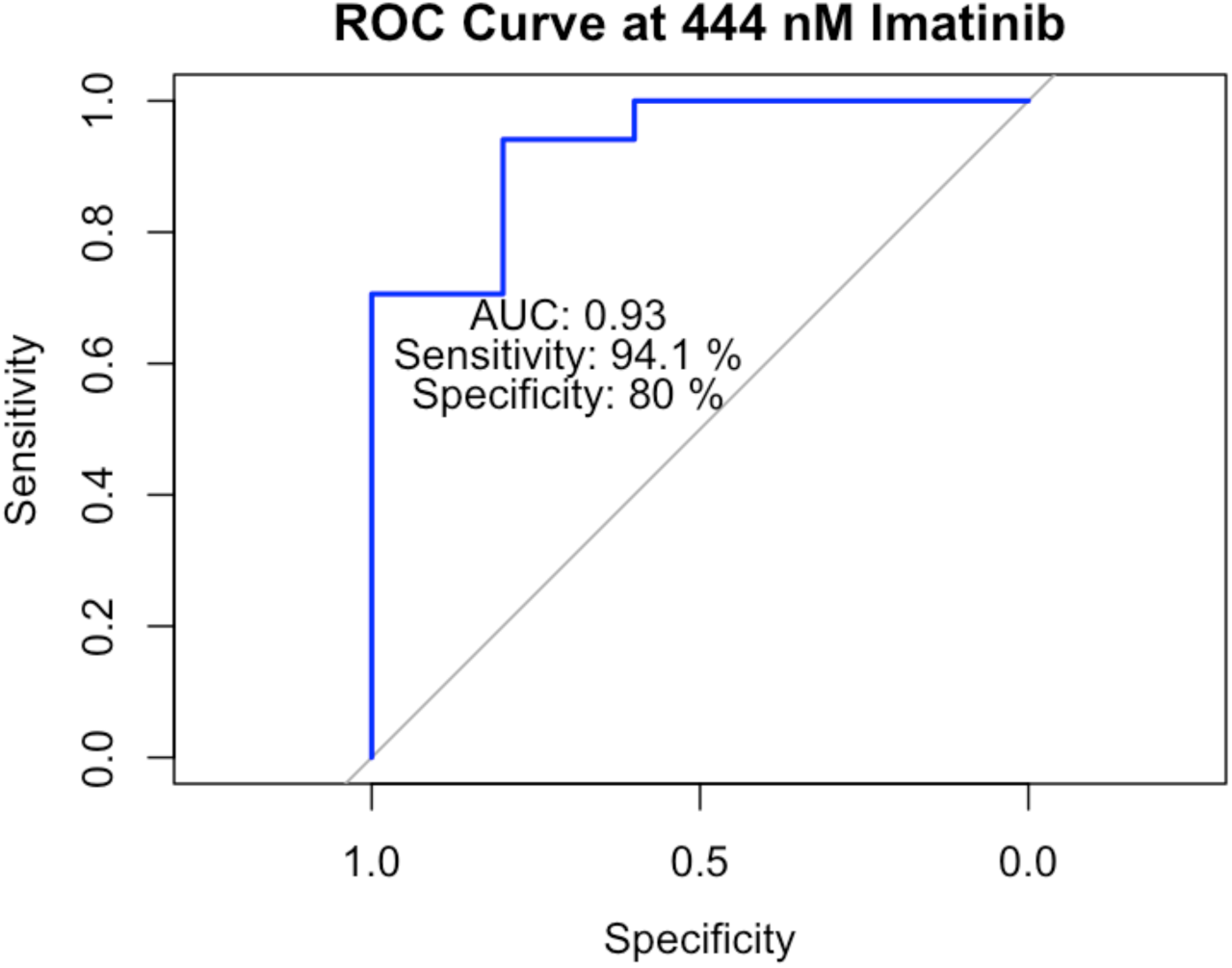
ROC curve shows the performance of the 4PL mutation classification model for discriminating resistant and sensitive single amino acid substitutions.

**Supplementary Figure 9.**
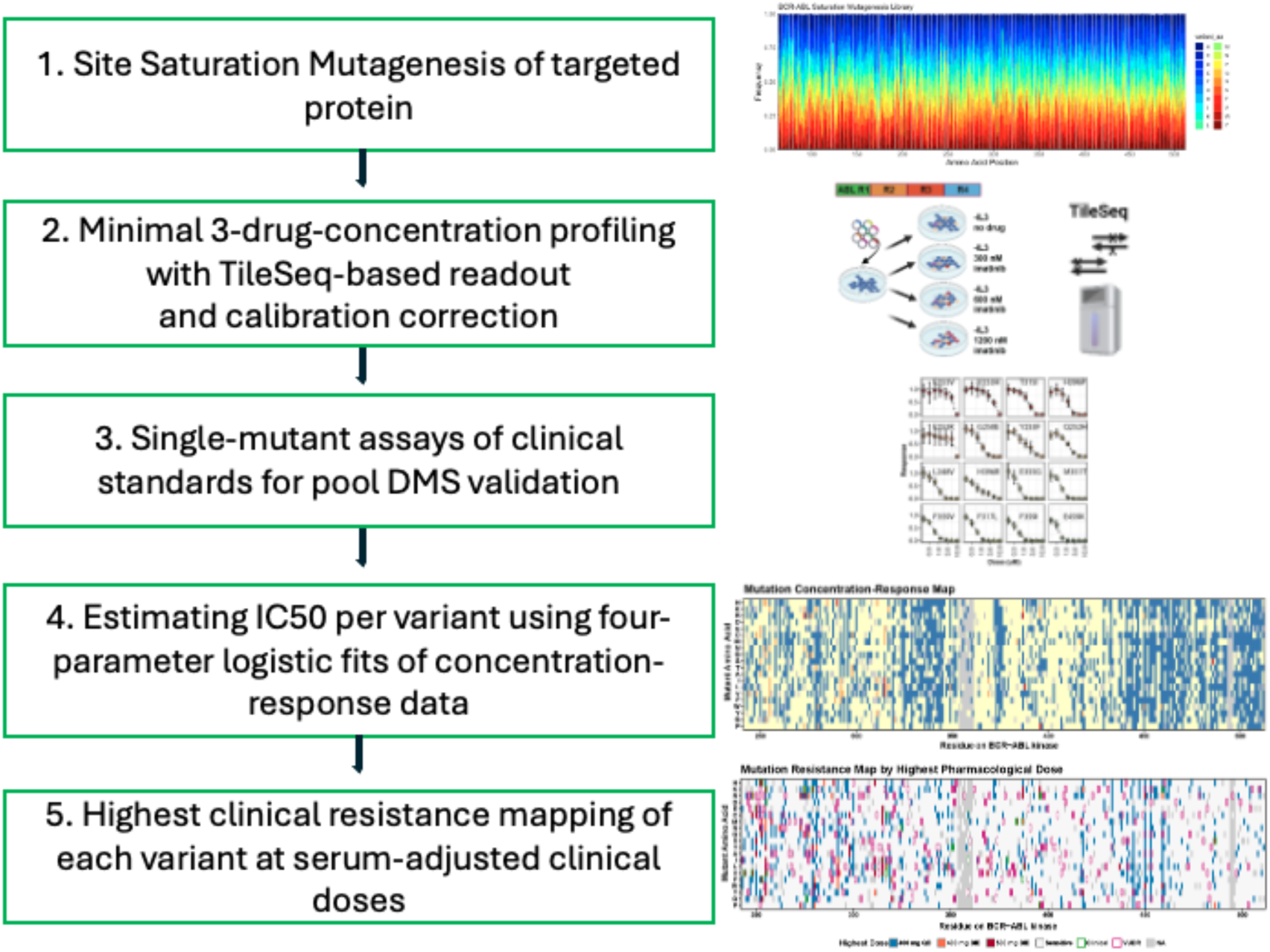
VUDR workflow for classifying clinical drug resistance of protein variants-specifically applied here to BCR-ABL variants - using a clinical dose-dependent deep mutational scanning (DMS) strategy integrated with PK/PD relationships. Step 1: The heatmap on the right visualizes library coverage across the protein’s amino acid sequence, indicating successful saturation mutagenesis.

